# Diffusion analysis of single particle trajectories in a Bayesian nonparametrics framework

**DOI:** 10.1101/704049

**Authors:** Rebeca Cardim Falcao, Daniel Coombs

## Abstract

Single particle tracking (SPT), where individual molecules are fluorescently labelled and followed over time, is an important tool that allows the spatiotemporal dynamics of subcellular biological systems to be studied at very fine temporal and spatial resolution. Mathematical models of particle motion are typically based on Brownian diffusion, reflecting the noisy environment that biomolecules typically inhabit. In order to study changes in particle behaviour within individual tracks, Hidden Markov models (HMM) featuring multiple diffusive states have been used as a descriptive tool for SPT data. However, such models are typically specified with an *a-priori* defined number of particle states and it has not been clear how such assumptions have affected their outcomes. Here, we propose a method for simultaneously inferring the number of diffusive states alongside the dynamic parameters governing particle motion. Our method is an infinite HMM (iHMM) within the general framework of Bayesian non-parametric models. We directly extend previous applications of these concepts in molecular biophysics to the SPT framework and propose and test an additional constraint with the goal of accelerating convergence and reducing computational time. We test our iHMM using simulated data and apply it to a previously-analyzed large SPT dataset for B cell receptor motion on the plasma membrane of B cells of the immune system.

## Introduction

Advances in fluorescence microscopic imaging have allowed us to develop an increasingly well-resolved picture of the spatial distribution and spatiotemporal mobility of important cellular proteins. Our rapidly developing knowledge has supported the development of quantitative theories of intercellular communication, cell surface receptor signaling and downstream responses.

Single particle tracking (SPT) is a technique that is of particular importance in defining the modes of protein mobility on the cell surface and within the cell [1]. Typically, cellular proteins are specifically labelled with a fluorescent tag. By labelling only a small fraction of molecules, individual fluorescent tags can be localized in a series of images and image analysis software can be used to link tag positions and thus obtain particle trajectories [2]. Once we have the trajectories, we can analyse their properties. Assuming that the particles are only subject to thermal noise, it is natural to analyze the tracks as representatives of simple Brownian motion and estimate the diffusion coefficient [3–5], while taking into account limitations in particle localization accuracy [6–9].

Particle trajectories extracted from cellular proteins have often exhibited deviations from simple diffusive behaviour, which has been attributed to transient binding to cellular structures, transient confinement in subcellular domains, directed motion under the influence of molecular motors, anomalous diffusion, etc [10–13]. In order to analyze transient behaviour within individual tracks, dynamic multi-state models have been developed and analyzed over the last ten years [14–19]. These methods explicitly assume that particles can switch among a specific number of diffusive states. For example, Das et al. used a two-state Hidden Markov Model (HMM) to fit particle tracking data for the surface receptor LFA-1 on the surface of T cells [14, 15]. In this model, particles were able to transition between two diffusive states, each characterized by a particular diffusion constant. The diffusion constants and transition rates between the states were fit to the data. At the time, previous experimental evidence pointed to a large reduction in LFA-1 mobility upon binding with cytoskeletal components, suggesting that the model could be a reasonable approximation to the dynamics of LFA-1 as it transitioned between bound and unbound states. More refined HMM approaches have subsequently been developed, reflecting finite localization errors and including particle capture within potential wells [20–22].

However, in the more general setting, how can one *a-priori* ascertain the best number of states that suffice to explain the data and provide insights into the biology of the tracked particles? Besides being a biological question of great interest, the number of possible states changes the statistical model and hence, the number of parameters to estimate. Therefore, estimating the number of states is an important problem in multi-state analysis.

One solution to this problem was recently introduced by Linden and Elf [23], where a variational Bayesian approximation was used to select the best model and a predictive approach was implemented using cross-validation. An alternative approach was previously described by Koo et al [24], where ensembles of short particle trajectories were analyzed through an expectation-maximization approach to a Gaussian mixture model, yielding the number of states, their diffusivities, and the stationary probabilities that particles inhabit each state. Here, we will attack the problem using a Bayesian nonparametric approach that allows the parameter space to be infinite-dimensional. The so-called Infinite Hidden Markov Model (iHMM) is a nonparametric model [25] that has recently been applied to FRET data by Pressé and coworkers [26, 27] to estimate the number of conformations of a molecule and simultaneously infer kinetic parameters for each conformational state. We use these concepts to develop a novel tool to analyze single particle tracking data under the assumption that the trajectories follow a Markov chain, where each element of the chain is a diffusion process. We seek to infer the number of diffusive states, the transition rates among states and the diffusion coefficient defining each state from the available data.

In this paper we will begin by specifying the HMM model that we wish to apply to SPT data, and then follow the thorough exposition of Sgouralis et al. [26] to generalize to the infinite-dimensional (iHMM) setting. We show validation using simulated data and provide a technical improvement to the algorithm that assists with convergence for the problem at hand. Finally, we apply our method to real data from experiments using TIRF microscopy to visualize motion of surface receptors on the membrane of live B cells [28] and discuss biological implications and possible future directions for study.

## Methods

### Infinite Hidden Markov Model for SPT (iHMMSPT)

We assume that tracked particles can be described through a memoryless process as follows. Suppose that particles switch among *K* different states where switching only occurs at the observation times (frames). Each state *σ_k_* yields spatial steps, corresponding to observed steps of the track, drawn from a Gaussian distribution with mean 0 and precision *ν_σ_k__*, where *k* = 1,…, *K*. Therefore, the probability distribution, *F_σ_k__*, of displacements for each state is given by

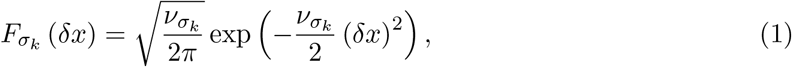

where *δx* is the displacement of the particle.

Experimental measurements give us the trajectory of each particle and thus the sequence of displacements, *δx_n_*, for each particle, *n* = 1,…, *N*. However, each displacement might be drawn from any of the diffusive states *σ_k_*. Thus, the sequence of states for the Markov model of this set of displacements is *s* = {*s*_1_,…, *s_n_*,… *s_N_*}, where *s_n_* might be any of the *σ_k_*. We can write this model in shorthand as

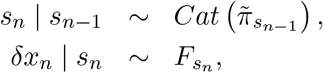

where 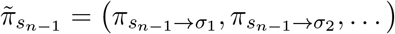 is a vector whose elements are the transition probabilities from the state *s*_*n*−1_ to any state *σ_k_*, and 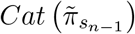 is the categorical probability distribution, whose weights are the transition probabilities. To estimate the parameters using HMM, we maximize the likelihood of observing the displacements given the model over the parameter set of the model.

However, for the iHMM case, we do not have a fixed number of states *K*. Therefore, we need to estimate the number of states *K*, the precisions *ν_σ_k__* and the transition probabilities. The iHMM model can be summarized as follows: we start with a number of states *K*, and use this to sample the transition probabilities. Next, we can use this information to sample the sequence of states, and subsequently the precisions *ν_σ_k__* using the emission model. The realization of the model can be written as follows:

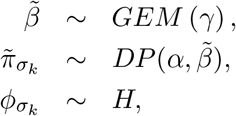

where *GEM*(*γ*) and 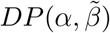 are the Griffiths-Engen-McCloskey and Dirichlet processes with concentration parameters *γ, α* and base distribution 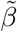 respectively. The prior of the parameters is *H*. For the SPT data, we use the conjugate model given by a Gaussian distribution with known mean, *μ_σ_k__* = 0, and unknown precision, *ν_σ_k__*, whose prior is a Gamma distribution [29]. Finally, in each realization, we use the data to learn the states and their respective parameters until convergence.

The Dirichlet process, 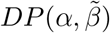, is a stochastic process whose range is a set of probability distributions. Analogously to sampling a familiar probability distribution, where we obtain a value for a random variable, in the Dirichlet process we sample probability distributions around the base distribution 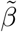, where *α* is a scaling parameter. For *α* ≪ 1, the *π_σ_k__* values are more likely to differ, while for *α* » 1, the *π_σ_k__* values are more likely to be similar.

We use the Griffiths-Engen-McCloskey process to generate the base probability distribution 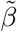 for the Dirichlet process. This is realized through the stick-breaking construction. It is defined as:

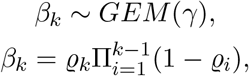

where *ϱ* are independent random variables drawn from a *β*(1, *γ*) distribution. The intuition behind this process is as follows: Imagine that one has a stick of unit length, which is then broken at a point *ϱ*_1_ drawn from *β*(1, *γ*). Then using the rest of the stick (of length 1 – *ϱ*_1_), select a new break-point *ϱ*_2_ = *ϱ*_2_(1 – *ϱ*_1_). Continuing this sequence of tasks, one ends up with variables *ϱ_i_* that have the two required probabilistic properties: they are each less than or equal to one, while their summation is equal to one, the length of the original stick. However, this distribution must be truncated because the number of states is potentially very large. We use the beam sampler to achieve this truncation. The beam sampler has an auxiliary variable that truncates the number of states. Let 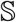 be the state space, where its size is equal to *K*. The auxiliary variable allows for an expansion in 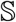 only when needed.

Let’s now define the auxiliary variable to be *u_i_*, for each time point *i* = 1 ··· *N* of a trajectory. The *u_i_* are drawn from a uniform distribution defined as: 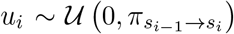. Then, the probability of a transition from state *σ_k_* in 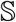 for any state outside 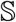 is equal to

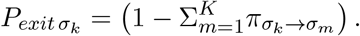

If the condition 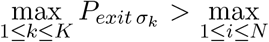 *u_i_* holds, then we do not have all the necessary states in the state space 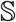 to explain the data, since the maximum probability of leaving the state space is larger than the maximum probability of transitioning between states inside the set 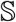. Thus, in this case we must add a new state in 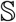, whose size will increase to *K* + 1 states. States will be added to 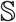 until the condition fails.

After the state space has been defined, the sequence of states *s* = {*s_i_*|*i* = 1,…, *N*} is sampled by using a forward-filtering backward sampling, which is a slight modification of the forwardbackward algorithm. Next, with the sequence of states for the trajectory, we can check whether there are any state of the state space 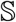 that has not been visited. Then, any state that has not been visited is deleted from the state space 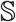. This step is called compression of space state. In summary, we expand the state space to avoid underfitting, and we compress 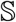 to avoid overfitting. Finally, we resample the parameters 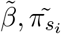, and the emission parameters: *ν_σ_k__*. There are many more technicalities in the general method that one can find in [26].

Finally, after obtaining the model and the parameters, one might be interested in estimaing the state that the particle occupies at each time point. To do this, we use the forward-backward algorithm to calculate the likelihood of being in state *σ_k_* given the number of states, their parameters, and the observations. We then select the state that gives the maximum likelihood in that specific time point [30].

### Improving convergence by accelerating state space compression

In the iHMM algorithm as described above, the compression of state space is a step that discards unnecessary states. After sampling the sequence of states, any state that has not been visited is discarded. Since they have not been visited, they did not produce any of the observations.

However, we propose that it is helpful to be more strict with the SPT data. For single particle tracks model, each diffusive state follows a Gaussian process with mean zero. This generates a lot of overlap among the distributions of each state. Thus, the creation of new states with precision close to the real one becomes common, and this slows convergence. This is illustrated in Figure 1. In this figure, we first plot the Gaussian distribution for each state for a model with *K_real_* = 4 states. We then iterated the iHMMSPT algorithm using the usual compression method and plotted the estimated Gaussian distribution for each state after 2000, 2500, 3000 and 3500 iterations (Fig. 1b). We can see that even after 3500 iterations, we do not achieve the correct number of states. However, if we increase the number of iterations from 3500 to 100000, the algorithm does converge (Fig. 2).

**Figure 1:**
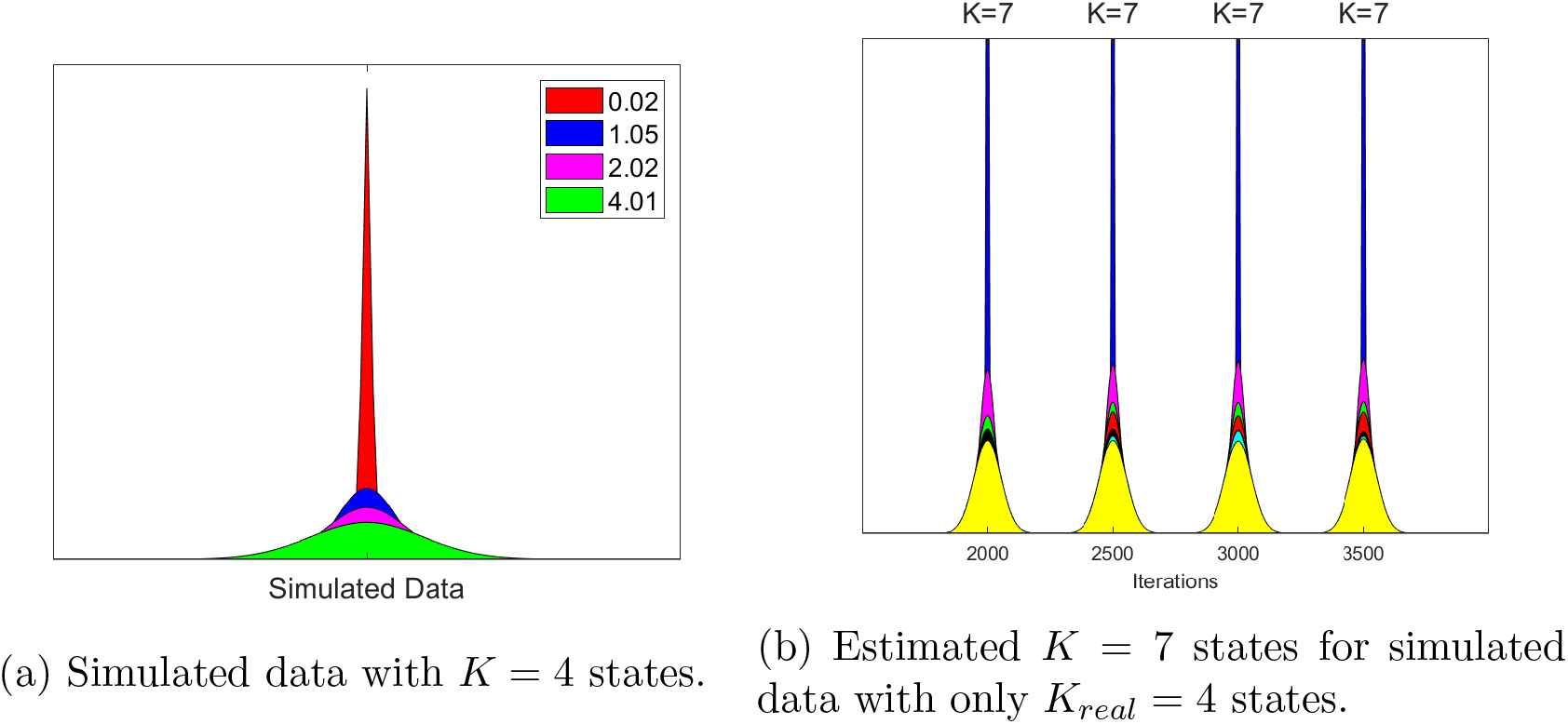
Graphs of the Gaussian distribution for each state: (a) for simulated data with *K_real_* = 4 states with diffusion coefficients as indicated, and (b) of estimated states using iHMMPST over 3500 iterations. The estimated number of states is *K* = 7, and we can see a lot of overlap among the Gaussians distributions of each state.

**Figure 2:**
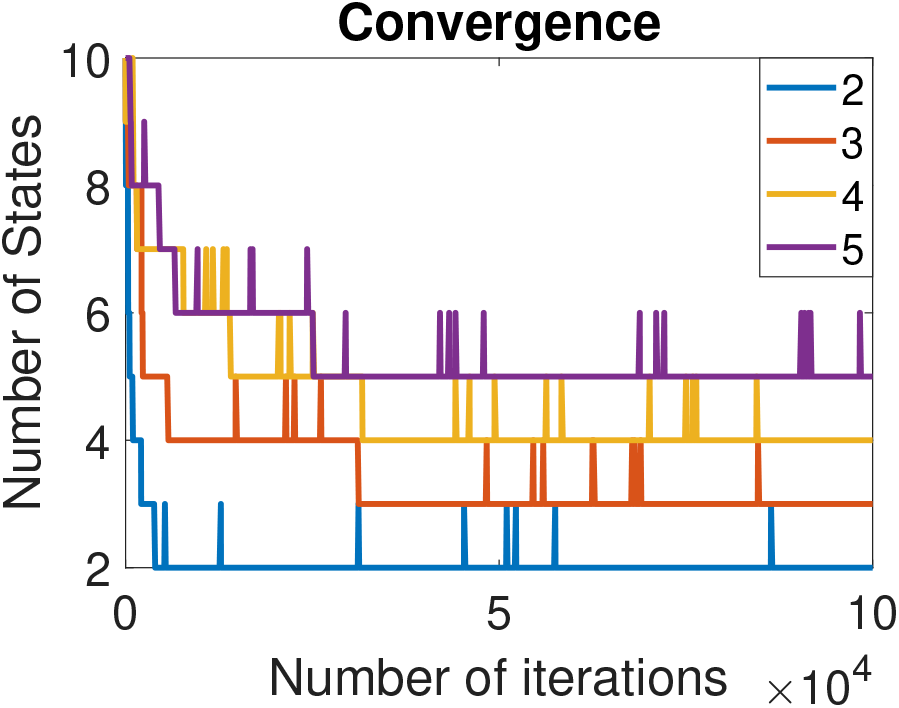
Convergence of algorithm without any additional conditions in the compression of states step. We ran the algorithm for 4 different sets of simulated data (2, 3, 4 and 5 states). We can see that the algorithm converges to the right number of states for all cases, after around 4 × 10^4^ iterations.

Seeking to accelerate convergence, we decided to add another condition to additionally compress the state space. We use the Bhattacharyya distance (Bd), a quantity that measures the similarity between two probability distributions, to decide if two states should be merged into one. For two Gaussian probability distributions with means equal to zero and precisions *ν*_1_, *ν*_2_, the Bhattacharyya distance is equal to

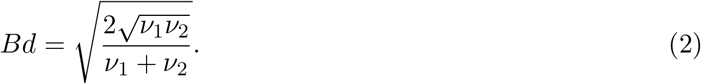

For equal precision, *ν*_1_ = *ν*_2_, we have *Bd* = 1, and as *ν*_1_ and *ν*_2_ grow apart, *Bd* → 0. We plot *Bd* as a function of the precision of two Gaussian distributions in Figure 3. We observe that *Bd* is higher than ~ 0.5 for the majority of the domain. In the iHMM algorithm, we will merge two different states into one if their pairwise *Bd* exceeds a given threshold. For simulated data, the threshold we use is 1 – 10^−6^. Since simulated data has a perfect accuracy, we choose a very large threshold to make sure no valuable information is lost.

**Figure 3:**
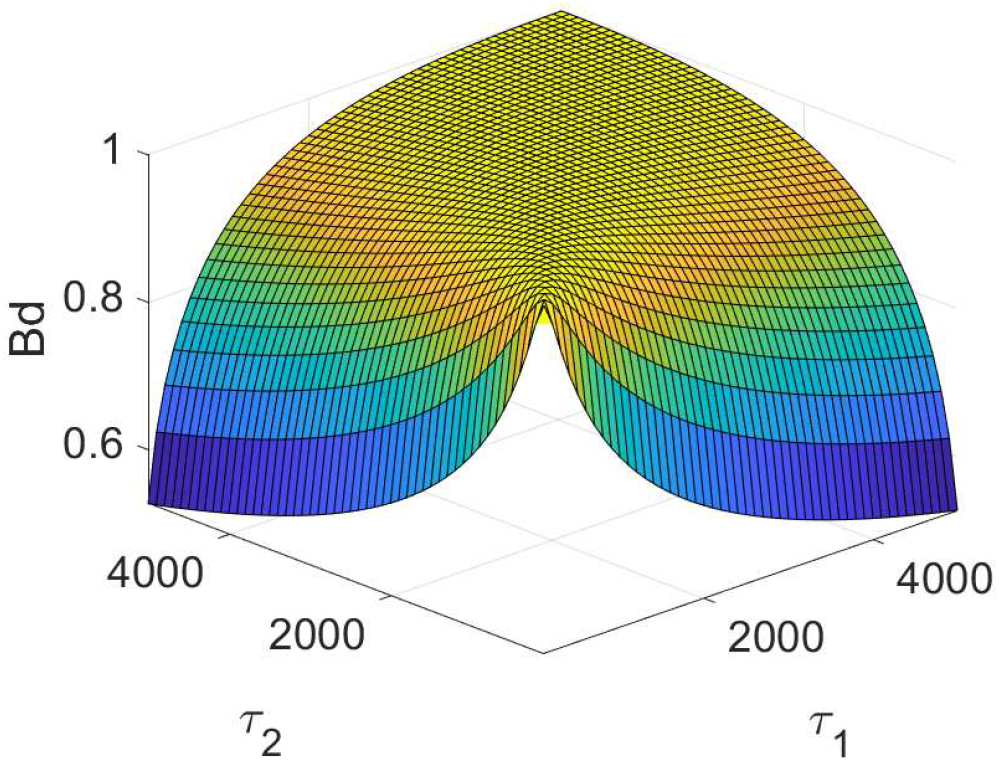
Bhattacharyya distance between two Gaussian distributions with the same mean but distinct precisions *τ*_1_ and *τ*_2_.

To determine the threshold for experimental data, we use a heuristic based on the localization accuracy of the data, as follows. Let *ε* be the measured localization accuracy of the experiment. We want to estimate a lower bound for the *Bd* threshold so that the differences between estimated diffusions exceed the accuracy of our experiment. Let *ν*_1_, and *ν*_2_ be the precisions oftwo states. If the difference between the diffusion coefficients of these states is within the experiment accuracy, then the diffusion coefficient of the second state has to be 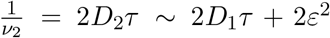, with 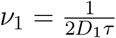. Therefore, the pairwise *Bd* for states with difference in the diffusion coefficients within the accuracy of the experiment, *Bd_acc_*, is given by:

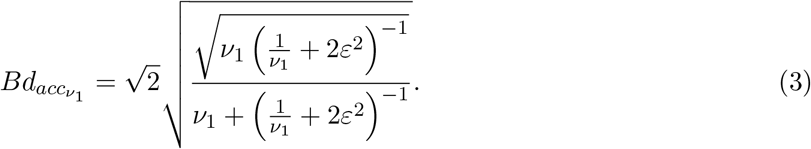

Using equation 3, we define a dynamical threshold for the experimental data. For each pair of states, (*m, n*), we calculate *Bd_acc_ν_m___* and *Bd_acc_ν_n___*. Therefore, the threshold for this pair of states, (*m, n*), is given by max(*Bd_acc_ν_m___, Bd_acc_ν_n___*).

## Results

### Algorithm testing with simulated data

We first simulate trajectories of particles whose motion is defined by steps drawn from a set of possible diffusive states. Particles transition between states with fixed rates, forming a Markov process. We then analyse the simulated trajectories with the iHMMSPT algorithm, to obtain estimates of the number of states, the diffusion coefficient of each state and the transition matrix of the Markov Model. Table 1 summarizes results for five example datasets, each with a different number of states. Each simulated particle is simulated in two dimensions, over 5 × 10^4^ frames, with a simulated frame-rate of 1000 frames per second.

**Table 1:** Results from testing the iHMMSPT algorithm with simulated data. We simulated 5 different datasets with 1 – 5 diffusive states. Here, we report the mode of the estimated number of states after burn-in (K), the mean estimated diffusion coefficient for each distribution (D), and the mean stationary probability (SP). The stationary probability is calculated from the estimated transition rate matrix.

| Simulated |  |  | Estimated |  |  |
| --- | --- | --- | --- | --- | --- |
| # of states | D | SP | K | D | SP |
| **5** | 0.0088 | 0.2279 | **5** | 0.0148 | 0.2432 |
|  | 1.0513 | 0.1744 |  | 1.2403 | 0.1122 |
|  | 2.0392 | 0.3257 |  | 1.9284 | 0.3917 |
|  | 4.0348 | 0.1513 |  | 4.1753 | 0.0998 |
|  | 8.0542 | 0.1207 |  | 7.5849 | 0.1531 |
| **4** | 0.0243 | 0.1344 | **4** | 0.0437 | 0.1595 |
|  | 1.0532 | 0.2457 |  | 1.1231 | 0.1150 |
|  | 2.0187 | 0.3403 |  | 1.8644 | 0.4741 |
|  | 4.013 | 0.2795 |  | 4.1065 | 0.2514 |
| **3** | 0.0434 | 0.2811 | **3** | 0.0528 | 0.2997 |
|  | 1.0106 | 0.3454 |  | 1.2401 | 0.4657 |
|  | 2.0313 | 0.3734 |  | 2.2645 | 0.2346 |
| **2** | 0.0336 | 0.8812 | **2** | 0.0343 | 0.8795 |
|  | 1.0078 | 0.1188 |  | 1.0507 | 0.1205 |
| **1** | 0.0412 | 1 | **1** | 0.0419 | 1 |

The algorithm is divided in two phases: a burn-in phase where we iterate the algorithm 1000 (20%) times using the additional condition to compress the state space, followed by 4000 (80%) further iterations using the usual compression step (Methods). After burn-in, the iterations of the process provide estimated distributions for number of states, diffusion coefficients and state transition matrix. In Table 1, we show the mode of the estimated number of states, the mean of the distribution for each diffusion coefficient and the mean of the stationary probability for each inferred state.

In Figure 4, we show the convergence of the number of states for each example. We observe that the algorithm converges rapidly to the correct number of states for each dataset during the burn-in step (the initial condition for all runs was *K* = 10). The diffusion coefficients are also key parameters of the model. In Figure 5 we show the final distributions obtained for the diffusion coefficients for each simulated dataset. We observe that the estimated diffusion coefficients are in good agreement with those used for simulation. Moreover, as we expect, the variance of the final distribution is inversely correlated with the stationary probability of each state. State 4 of the 5 states dataset is a good example of this (see also Table 1).

**Figure 4:**
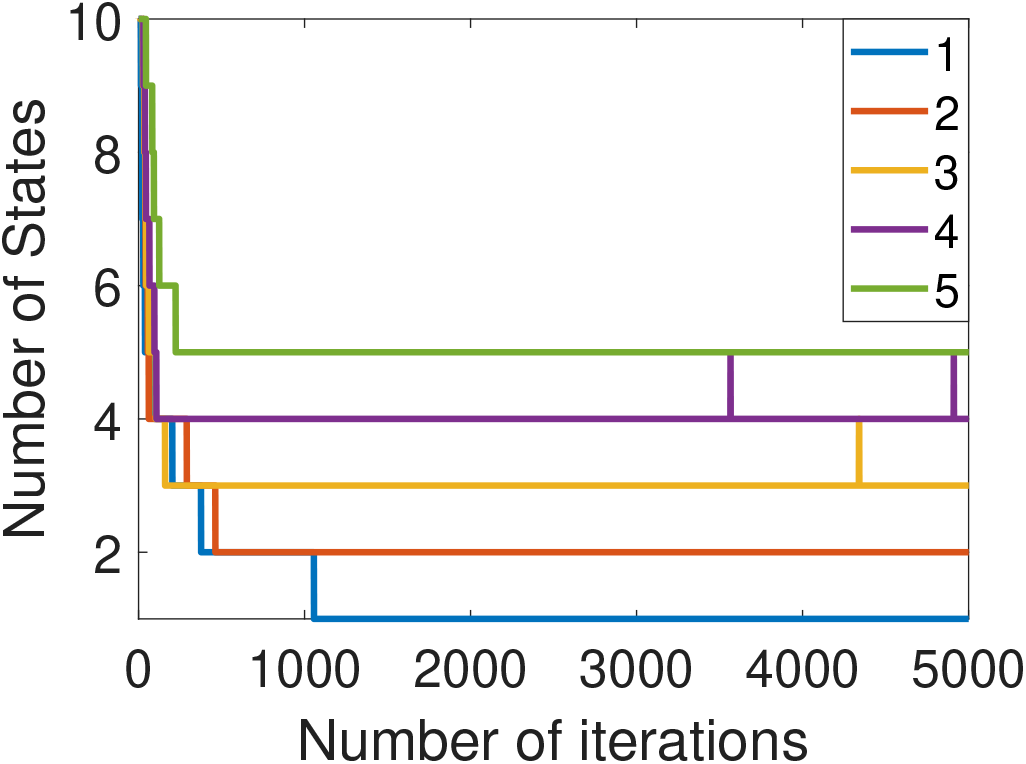
Plot of the number of states of each iteration for each simulated dataset using the additional compression criterion during burn-in. Simulations with one to five states were fitted. We can see that after about 1000 iterations we achieved good convergence for all cases.

**Figure 5:**
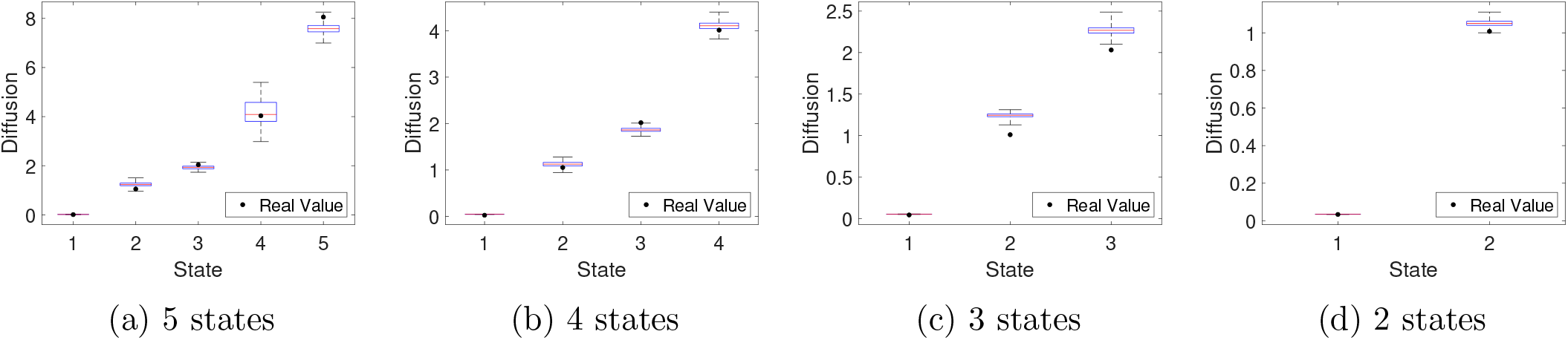
Boxplots of estimated diffusion coefficient distributions for simulated data with different numbers of states. Black circles represent true diffusion coefficients.

We observe from Table 1 that the algorithm estimates the correct (simulated) number of states K for every dataset. We also obtain generally good estimates of the diffusion coefficients. The signs of errors in estimation of diffusion coefficients are in line with our expectations. For example, for *K* = 4, we have two states with true diffusion coefficients equal to 1.0532 and 2.0187. The estimated diffusion coefficients are 1.123and 1.8644, respectively. These discrepancies occurred because some displacements that are actually from the slower states have been treated as displacements from the faster state. We can confirm this by looking at the stationary probabilities. The faster state stationary probability was overestimated, while the stationary probability of the slow state was misunderestimated. However, in general, we find that the algorithm performs well with simulated data of this type, even though the number of parameters to be estimated is quite large. For example, when *K* = 4, we have 20 parameters to estimate: 4 diffusion coefficients and 16 transition probabilities.

Finally, we performed state segmentation on each simulated trajectory by finding the most probable state at each frame, given the estimated parameters for that trajectory. We then compared the estimated state sequence with the true simulated state sequence and calculated the percentage of displacements that are correctly classified (Table 2). We observe that the accuracy of the state segmentation decreases as the number of states grows. This is expected because the quality of the diffusion and transition parameter estimates generally decreases as the number of states increases, and this affects the quality of state segmentation.

**Table 2:**
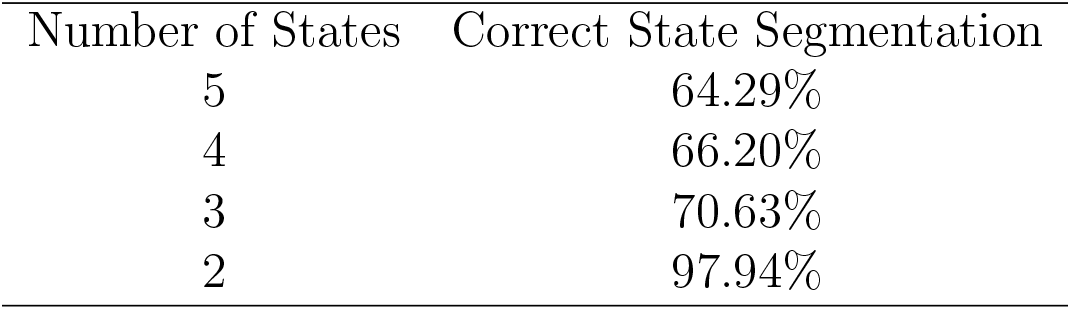
Accuracy of state segmentation from simulated trajectories.

| Number of States | Correct State Segmentation |
| --- | --- |
| 5 | 64.29% |
| 4 | 66.20% |
| 3 | 70.63% |
| 2 | 97.94% |

### Application to B cell receptor tracking data

We now apply the iHMMSPT algorithm to experimental data. In previous work, we performed a detailed comparison of results from SPT using two different methods for labeling cell-surface proteins. Briefly, B cell receptors (BCR) on the surface of B lymphocytes were labeled using quantum dots (Qdots) linked to monovalent antigen-binding fragments of antibodies (Fab), or directly conjugated to the small organic fluorophore Cy3. Images were taken at 30 frames per second using a total internal reflection fluorescence (TIRF) instrument. Tracks were extracted from image stacks using Icy software [31]. The spatial precision of particle localization was estimated to be 23nm and 30nm for Qdot and Cy3 labelling, respectively. Full methods are reported in our original paper [28].

Using this data, we previously conclusively showed that the mobility of proteins labeled using Qdots was impaired, most probably due to steric hindrance. As part of our analysis, we used a two-state HMM to segment the particle tracks among two diffusive states: a fast and a slow state [14]. Using the iHMMSPT algorithm, we can now answer the question as to whether the data is better described by a different number of states. This approach has the potential to give us insights into the heterogeneity of potential interactions between the tracked receptor and different systems at the cellular membrane, such as the cortical actin cytoskeleton, lipid rafts, transiently binding proteins, etc. We can also compare between the two labelling strategies and obtain a refined picture of changes in the diffusive behaviour.

We examine six datasets, each from a separate experiment. Three experimental datasets used directly-conjugated Cy3-labelling of IgG and three used Qdot-labelling of IgG [28]. For the Cy3 experiments, the datasets contained 1886, 1768 and 1733 tracks, respectively, with average numbers of frames per track of 58 (standard deviation 77). For the Qdot experiments, the datasets contained 2394, 4468 and 1526 tracks, respectively, with average numbers of frames per track of 83 (standard deviation 94). Before applying our iHMMSPT algorithm, we first applied an immobilility threshold to remove immobile tracks, as previously described [28]. We allow the algorithm to complete 10000 iterations in total, with a burn-in phase of 3000 iterations. For each trial, we run the algorithm with the additional *Bd* condition chosen based on localization accuracy (Methods).

Results are shown in Tables 3 and 4. The diffusion coefficients and stationary probabilities reported are the mean of the distributions of each state of all iterations. We also report the estimated full transition matrices for each labelling method in the Appendix.

**Table 3:** iHMMSPT results from three sets of experimental tra jectories obtained from IgG molecules directly conjugated to Cy3 on the surface of live B cells. For each experiment, we acquire 5 states as the optimal number of states. Mean estimated diffusion coefficients (D) and stationary probabilities (SP) are also reported for each state.

| ***Labelling*** | Cy3 1 |  | Cy3 2 |  | Cy3 3 |  |
| --- | --- | --- | --- | --- | --- | --- |
| *Parameters* | D($\mu\text{m}^2/\text{s}$) | SP | D($\mu\text{m}^2/\text{s}$) | SP | D($\mu\text{m}^2/\text{s}$) | SP |
| **1** | $< 10^{-5}$ | 0.1289 | $< 10^{-5}$ | 0.1321 | $< 10^{-5}$ | 0.1320 |
| **2** | 0.0061 | 0.0440 | 0.0094 | 0.0608 | 0.0119 | 0.0786 |
| **3** | 0.0259 | 0.2893 | 0.0375 | 0.3218 | 0.0421 | 0.2892 |
| **4** | 0.1060 | 0.4745 | 0.1337 | 0.4384 | 0.1545 | 0.4424 |
| **5** | 0.6625 | 0.0633 | 0.7510 | 0.0468 | 0.8303 | 0.0578 |

**Table 4:** iHMMSPT results from three sets of experimental trajectories obtained from IgG molecules directly conjugated to Cy3 on the surface of live B cells. Experiments 1 and 3 were found to support five states, while experiment 2 supported four states. Mean estimated diffusion coefficients (D) and stationary probabilities (SP) are also reported for each state.

| ***Labelling*** | Qdot 1 |  | Qdot 2 |  | Qdot 3 |  |
| --- | --- | --- | --- | --- | --- | --- |
| *Parameters* | D($\mu\text{m}^2/\text{s}$) | Prob | D($\mu\text{m}^2/\text{s}$) | Prob | D($\mu\text{m}^2/\text{s}$) | Prob |
| **1** | $< 10^{-5}$ | 0.1544 | $< 10^{-5}$ | 0.1863 | $< 10^{-5}$ | 0.1537 |
| **2** | 0.0053 | 0.1966 | 0.0073 | 0.452 | 0.0047 | 0.1757 |
| **3** | 0.0217 | 0.39311 | 0.027 | 0.3201 | 0.0181 | 0.4485 |
| **4** | 0.0766 | 0.2297 | 0.2785 | 0.0415 | 0.0582 | 0.2036 |
| **5** | 0.5470 | 0.0262 |  |  | 0.3760 | 0.0186 |

Interestingly, we find that the optimal number of states is five for all experiments, with both labelling methods, except for the second trial of Qdot-labelling experiment, whose optimal number of states is found to be four. We also find that the parameter estimates for each state are qualitatively similar from day to day. Examining the states in detail, we see that the first state is very slow for all experiments, reflecting particles that are transiently in an immobile state (recall that entirely immobile particles are removed from the data before analysis). Meanwhile, the largest diffusion coefficients (fifth state) are very large, estimated to be on the order of 0.1 – 1 *μ*m^2^s^−1^. The stationary probabilities for these show that they account for 5-6% of all states for Cy3-labelled receptors, but only 2-3% for Qdot-labelled receptors. Since Qdots are brighter and less prone to blinking than Cy3 molecules, this suggests that some or all of these larger steps are in fact due to tracking errors. We also find three intermediate mobile states. For all these, the diffusion coefficients estimated from the Cy3 experiments are a little larger than the diffusion constants for the respective states of the Qdot experiments, indicating that protein mobility was generally impaired due to Qdot labelling. Moreover, the most-occupied state for the Cy3-labelled molecules is the fourth state (*D* ~ 10^-1^*μ*m^2^s^−1^), whereas for the Qdot-labelled molecules, the most-occupied state is the slower third state (*D* ~ 10^-2^*μ*m^2^s^−1^). For the second trial of Qdot-labelled molecules, where we have 4 states, the most-occupied state is the second state (*D* ~ 5 × 10^-3^*μ*m^2^s^−1^). We also calculated the effective diffusion coefficient defined as *D_eff_* = *P*_1_*D*_1_ + *P*_2_*D*_2_ + ··· + *P_n_D_n_*. We obtain similar results to our previous work [28] for all datasets (not shown). Overall, our results are consistent with the hypothesis that the Qdot-labelled receptors can easily become trapped in small regions of the cell membrane, while the Cy3-labelled molecules can escape these regions and explore the cell membrane.

In Figure 6, we plot trajectories from two experiments, segmented (colour-coded) by diffusive state. Most Qdot trajectories are observed to be barely mobile and highly localized in very small regions, and mostly in the third state (red). On the other hand, the Cy3-labelled receptors are mostly in their (faster) fourth state (cyan) and are able to explore the cell surface. We can also observe transitions between states over time.

**Figure 6:**
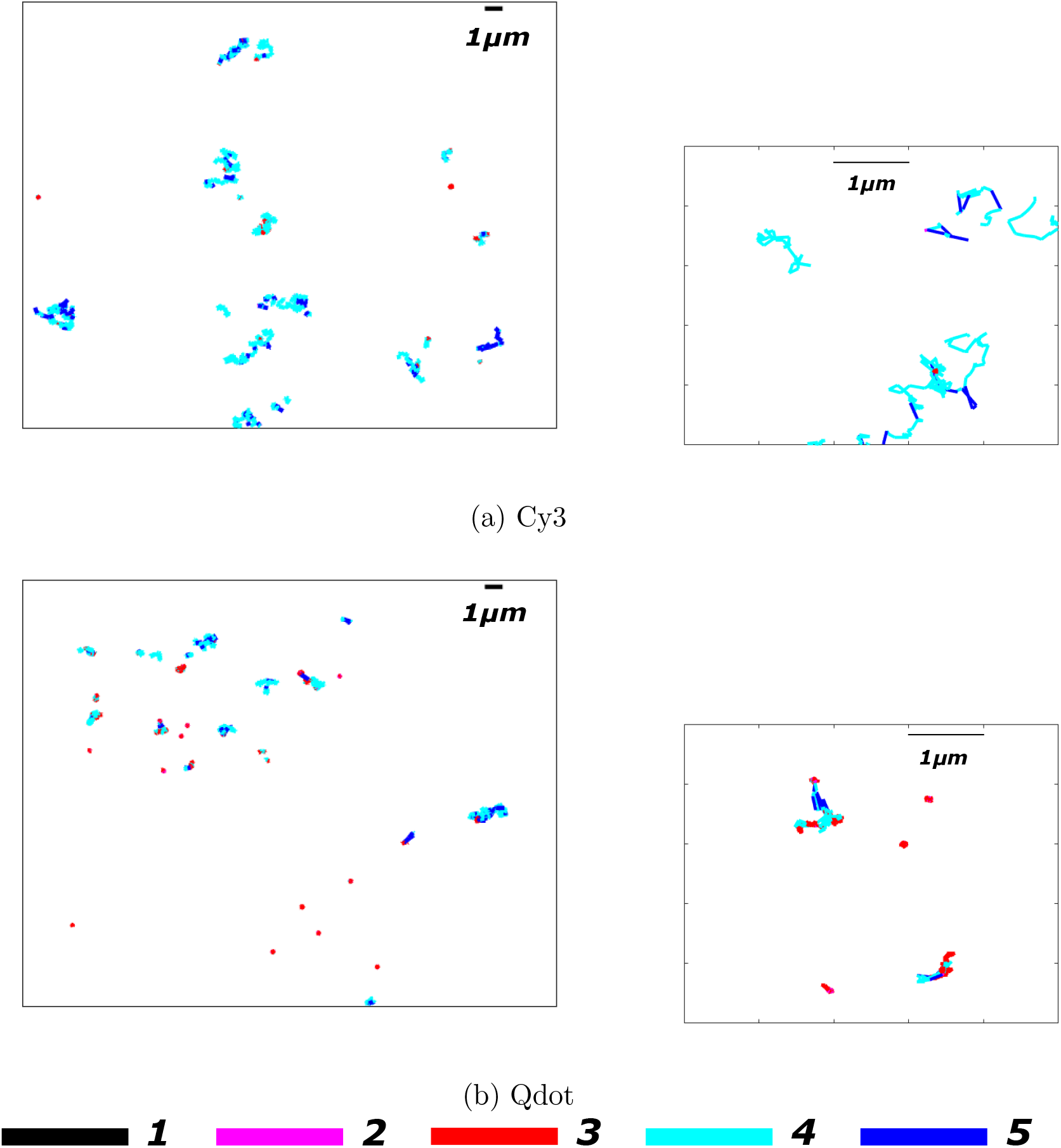
Segmented trajectories of (a) Cy3-labelled B cell receptors, and (b) Qdot-labelled B cell receptors, with a zoomed image of a smaller region. Colours represent the different diffusive states of each experiment. Diffusion constant estimates are presented in Tables 3 and 4. The trajectories of Qdot-labelled molecules are spatially restricted compared to those of Cy3-labelled molecules. Moreover, the Cy3 trajectories are estimated to frequently exist in the more-mobile states (*D* ~ 10^−1^*μ*m^2^s^−1^), whereas the Qdot trajectories are mostly found in slower states (*D* ≲ 10^−2^*μ*m^2^s^−1^).

These results about the impaired mobility of Qdot-labelled receptors are in line with previous work [28]. However, we previously imposed a two-state model, while here we find that five states are optimal to describe the data. Even after possibly discarding the fastest inferred state as likely due to tracking errors, we find that receptor mobility is more heterogeneous than previously imagined. In general, using a model with only two states is likely to cause the loss of potentially important information.

### Effects of cortical actin cytoskeleton disruption on BCR mobility

As a second case study, we re-examined a second set of data where the submembrane actin cytoskeleton of the B cell was disrupted by latrunculin A (LatA). Two experimental datasets used Cy3-labelling on IgM BCRs, and were treated with either DMSO (control) or LatA, and another two experimental datasets used Qdot-labelling on IgM BCRs and were treated with either DMSO (control) or LatA [28]. For the Cy3 experiments the datasets contained 3281 and 2963 tracks, respectively, with average numbers of frames per track of 50 (standard deviation 60). For the Qdot experiments the datasets contained 9262 and 8857 tracks, respectively, with average numbers of frames per track of 57 (standard deviation 60).

In all experiments, we found very slow (*D* ~ 10^−7^*μ*m^2^s^−1^) and very fast (*D* ≥ 1.1*μ*m^2^s^−1^) diffusive states. Again, the slow state shows transient confinement, while the fast state probably reflects a certain fraction of tracking errors. Also as before, the fastest state was more heavily weighted in the Cy3 experiments, probably reflecting a larger number of tracking errors compared to the Qdot experiments. Our results also show that labelled IgM is generally more mobile than labelled IgG, in agreement with previous analysis [28].

We did not find any major differences between control and LatA conditions when using Qdot labels. This tells us that the actin cytoskeleton is not the primary constraining factor; we believe that the steric hindrance of the large Qdot label is much more important. For the Cy3 label experiments, we find interesting differences between control and LatA. In agreement with previous analysis, receptors on LatA-treated cells are generally much more mobile than on control cells, implicating the actin cytoskeleton as an important regulator of receptor mobility. Comparing results between control and LatA treatment, we find that the most-occupied states for the control cells is the third state (*D* ~ 5 × 10^−2^*μ*m^2^s^−1^ in both cases), whereas for LatA, it is the fourth state (*D* ~ 1.8 × 10^−1^*μ*m^2^s^−1^ in both cases), indicating an increase in receptor mobility via a transition to a more rapid state. Compared to two-state analysis, we obtain a refined picture of the heterogeneity of the system.

## Discussion

In this paper we have described and implemented a novel iHMM algorithm for the problem of multiple state discrimination for SPT data analysis. We have also demonstrated its use on a collection of high-quality data that was previously analyzed using a fixed two-state HMM. We believe that the iHMM approach is a rational choice for this longstanding state counting problem and is superior to alternative approaches such as the use of information criteria to distinguish among multiple models. We were able to also improve the convergence of the algorithm via a simple approach of merging nearby states.

We used our algorithm to re-examine SPT data obtained for BCRs moving on the surface of live B cells, using two different labelling strategies and with pharmacological perturbation of the membrane actin cytoskeleton. We discovered evidence that BCRs can transition among approximately five distinct diffusive states, with a wide range of diffusion coefficients. The slowest detected states appear to reflect transient immobility, while the fastest state probably reflects occasional tracking errors. Our results on the difference between Qdot-labelled molecules and Cy3-labelled molecules are in general agreement with our previous work [28], where we conclude that Qdot-labelling impairs molecule mobility, but present intriguing possibilities for further study to understand the significance of the intermediate states in biological terms. Moreover, our analysis confirms previous results of the actin cytoskeleton being an important receptor regulator. On LatA-treated cells, we observe higher mobility of the receptors, suggesting that disruption of actin cytoskeleton promotes faster BCR motion.

In recent work, Rey-Suarez et al. [32] found that BCRs exist in 8 distinctive diffusive states. They used an expectation maximization approach based on a Gaussian mixture models, and allowing no transitions between states. In their work, the trajectories were split into segments each containing 15 frames. Each segment was assumed toarisefromasingle diffusive state. Similar to our findings here, they report a large number of diffusive states, confirming the heterogeneity and complexity of the cell membrane. However, since no transitions are allowed within each 15-frame segment, some segments could arise from a combination of diffusive states, yielding an overestimate of the optimal number of states. Nonetheless, the work of Rey-Suarez supports the use of multi-state models and shows their potential to support biological discovery.

Our approach to this problem has some potential weaknesses that we intend to address in future work. We have applied a restricted model of particle motion as a multi-state diffusion process. An alternative line of attack could allow for transient confinement of the particle within a potential well, or other forms of nondiffusive motion, as has previously been implemented by others in HMM and other settings [20, 21, 33]. Modified HMM models reflecting these kinds of additional complexity should be relatively simple to implement within the iHMM framework. Second, we did not account for localization errors in our current implementation. This is known to reduce the accuracy of estimation of diffusion coefficients in related settings [6], and was incorporated into the recent study of Rey-Suarez et al [32]. Finally, we have not performed a thorough analysis of our algorithm’s performance on simulated data, where questions of model mis-specification, as well as heterogeneity among track populations (as might arise from cell-to-cell heterogeneity in a single experiment) can be precisely examined. We intend to address all these weaknesses in the near future.

In summary, we have presented a generalizable approach that extends the potential applicability of HMMs for SPT data. More broadly, we have shown a novel application of the iHMM method, further proving that it is an excellent tool for quantification of experimental biophysics data [26]. All experimental datasets and software are available from the authors on request.

**Table 5:** Results of iHMMSPT for experimental data on B cells, where IgM were labelled using either directly conjugated fab (Cy3) or Qdots. For these experiments, the cells were treated with either DMSO control or LatA. We find no meaningful difference between the Qdot DMSO and Qdot LatA experiments. We find that Cy3-LatA tra jectories show faster state-wise diffusion coefficients compared to control.

| ***Exp*** | Cy3 DMSO |  | Cy3 LatA |  |
| --- | --- | --- | --- | --- |
| *States* | D($\mu\text{m}^2/\text{s}$) | Prob | D($\mu\text{m}^2/\text{s}$) | Prob |
| **1** | $< 10^{-5}$ | 0.1172 | $< 10^{-5}$ | 0.1161 |
| **2** | 0.0136 | 0.1946 | 0.0118 | 0.0555 |
| **3** | 0.0560 | 0.3973 | 0.0383 | 0.2587 |
| **4** | 0.1747 | 0.2399 | 0.1841 | 0.4590 |
| **5** | 1.1049 | 0.0510 | 1.0801 | 0.1107 |
| ***Exp*** | Qdot DMSO |  | Qdot LatA |  |
| *States* | D($\mu\text{m}^2/\text{s}$) | Prob | D($\mu\text{m}^2/\text{s}$) | Prob |
| **1** | $< 10^{-5}$ | 0.1691 | $< 10^{-5}$ | 0.1630 |
| **2** | 0.0088 | 0.3218 | 0.0064 | 0.2207 |
| **3** | 0.0309 | 0.3872 | 0.0254 | 0.4054 |
| **4** | 0.1744 | 0.0880 | 0.1086 | 0.1653 |
| **5** | 1.5255 | 0.0339 | 1.2586 | 0.0457 |

## Acknowledgements

We thank Libin Abraham, Michael Gold, Alejandra Herrera, Michael Irvine, Joshua Scurll and especially Tommi Muller for valuable discussions. This work was supported by a Discovery Grant from the Natural Science and Engineering Research Council of Canada (RGPIN-2015-04611 to DC). DC gratefully acknowledges sabbatical funding from the Quantitative Theory and Methods initiative at Emory University. We also thank the organizers of “Reverse mathematical methods for reconstructing molecular dynamics in a single cell” held at the Centro di Ricerca Matematica Ennio De Giorgi, in October 2018, for a stimulating and enjoyable workshop.

## Appendix

In this appendix, we provide the full estimated transition matrices (*E*) for the examples presented above, for simulated and experimental data. For simulated data, we also provide the true transition matrix (*S*).

**Simulated data with two states:**

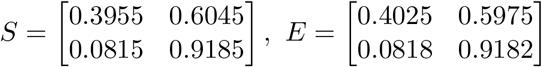

**Simulated data with three states:**

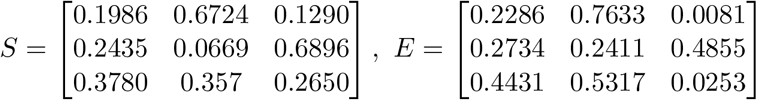

**Simulated data with four states:**

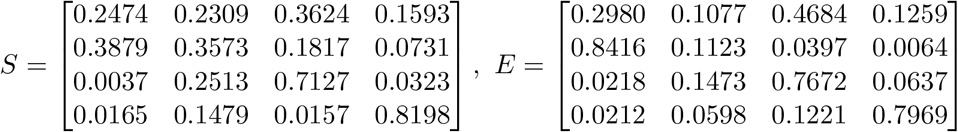

**Simulated data with five states:**

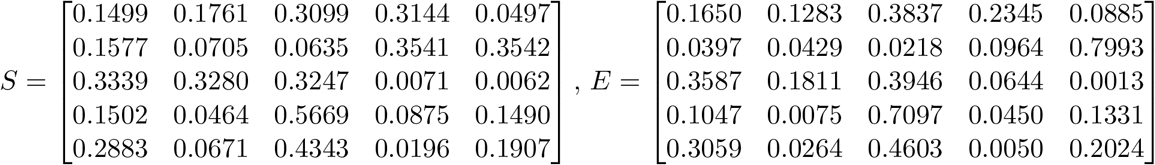

**Directly conjugated Fab (Cy3) on IgG experiment, trial 1 (Cy3 1):**

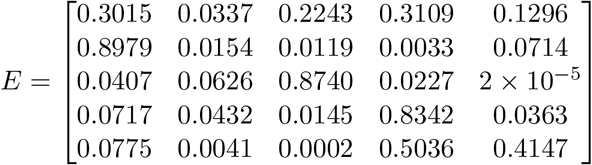

**Directly conjugated Fab (Cy3) on IgG experiment, trial 2 (Cy3 2):**

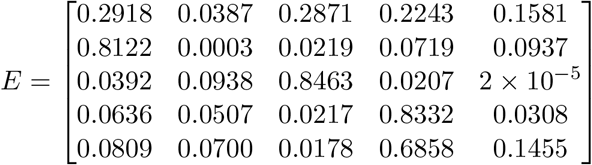

**Directly conjugated Fab (Cy3) on IgG experiment, trial 3 (Cy3 3):**

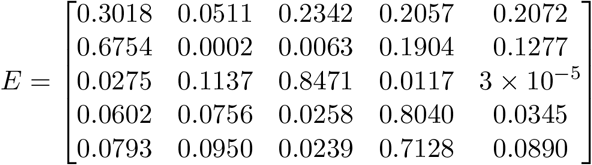

**Quantum dot experiment on IgG, trial 1 (Qdot 1):**

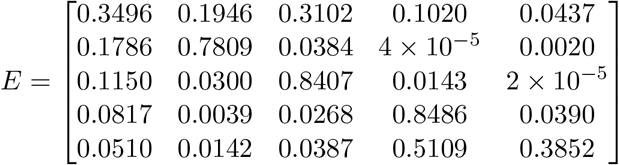

**Quantum dot experiment on IgG, trial 2 (Qdot 2):**

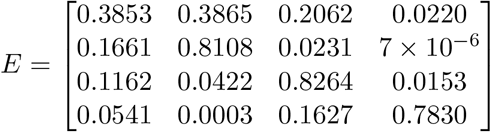

**Quantum dot experiment on IgG, trial 3 (Qdot 3):**

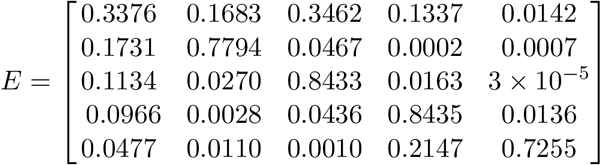

**Directly conjugated Fab (Cy3) on IgM + DMSO:**

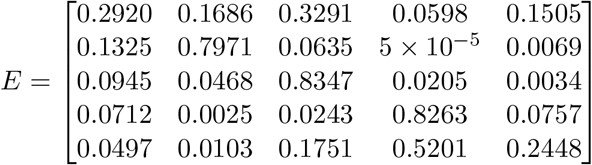

**Directly conjugated Fab (Cy3) on IgM + LatA:**

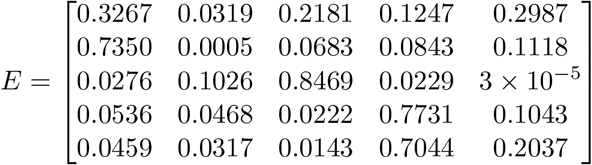

**Quantum dot experiment on IgM + DMSO:**

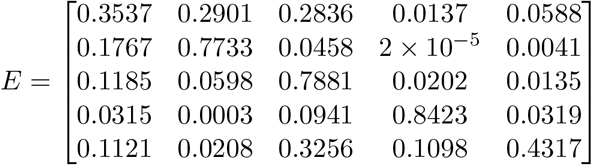

**Quantum dot experiment on IgM + LatA:**

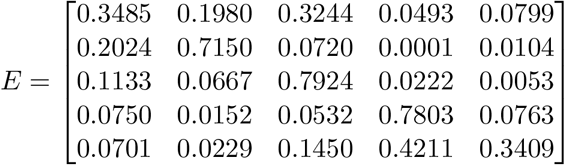

